# Evolutionary stability of frequency-dependent secondary defense in a soft-bodied insect with and without signals

**DOI:** 10.64898/2026.10.22.761097

**Authors:** Nich W Martin, Colette M St Mary

## Abstract

Secondary defenses can reduce predation risk by altering predator behavior, but their evolution may be constrained when defended prey are indistinguishable from undefended individuals. This is particularly relevant for soft-bodied prey, for which predator attacks are lethal regardless of defense level, such that the benefits of defense are frequency-dependent. We developed two sets of evolutionary models to investigate how secondary defense evolves in the presence vs. absence of signals that allow predators to distinguish defended from undefended prey. First, we used deterministic evolutionary game theory to examine the stability of defended and undefended strategies across attack likelihoods, defense levels, and initial frequencies of defended prey. Second, we developed stochastic, multi-generational models in which predators optimized foraging decisions according to their energetic state, allowing selection to emerge from state-dependent predator behavior. For both model sets, undefended prey were evolutionarily stable when predation pressure was low, whereas stronger predation favored defended phenotypes. However, in the absence of signals defended and undefended prey frequently coexisted, with stable mixed populations emerging when predation pressure was sufficiently high, defense levels low, and the initial proportion of defended prey was above a threshold frequency. In contrast, when signals were present, mixed equilibria were generally unstable and highly defended prey frequently displaced other phenotypes, particularly under high predation pressure. Stochastic models produced similar patterns while demonstrating that 1. prey population size influenced the likelihood of invasion by defended phenotypes, with smaller populations being more conducive to invasion; 2. qualitative patterns similar to the deterministic models emerged when selection was dependent on predators’ state-dependent behavior. Our results identify conditions under which inconspicuous, costly secondary defenses can persist even when defense provides no benefit to the individual, presenting a plausible route towards the evolution of phenomena such as aposematism.

## Introduction

Prey can decrease their likelihood of being captured and killed by predators through investment in defense. While some defenses operate by avoiding predator detection or attack (e.g., crypsis and warning signals, respectively) many defenses only become relevant once a predator has attempted to subjugate and/or consume an individual (e.g., toxins, spines, claws, etc.). The latter of these two defense types are often referred to as secondary defenses, while the former are considered primary defenses (Edmunds, 1974). Of course, primary and secondary defenses are not mutually exclusive (e.g., prey could be both cryptic and chemically defended). In many cases primary and secondary defenses can be synergistic. For example, warning signals (primary defense) are widely considered to facilitate learning among predators as to which prey are adopting secondary defenses and which are not, a phenomenon known as aposematism (Mappes et al., 2005).

While there is a vast literature of both empirical and theoretical work regarding the evolution of secondary defenses and warning signals, little work has been done with respect to the evolution of secondary defenses among inconspicuous prey indistinguishable from undefended members of their cohort (although see, Hansen et al., 2010; Svenningsen and Holen, 2007). This absence in the literature does not reflect that such strategies are rare, as there are a number of species utilizing secondary defenses lacking noticeable aposematic warning coloration (e.g., Koskenpato et al., 2026). Another reason studies of cryptic but secondarily defended prey are important is that theory predicts aposematic populations likely evolve from ancestral populations lacking secondary defenses via a transitional population state, where defended individuals are indistinguishable from undefended individuals (Ruxton et al., 2019). For such a state to evolve and persist, secondarily defended (hereafter simply, defended) prey must arise and increase in frequency in a population with undefended individuals. This begs the question, if predators are incapable of distinguishing defended prey from undefended prey, what conditions allow defended individuals to invade a population, and how do these differ from scenarios in which predators are capable of distinguishing between prey types?

Broom et al. (2005) modeled scenarios in which identical individuals with defenses invisible to predators vary along a defense level continuum. They showed that when defenses are continuous, populations are likely to converge on a single defense level. Svennungsen and Holen (2007) modeled a similar scenario in which defenses can be detectable, but with so-called “automimics” present in the population, i.e., identical undefended individuals. These automimics can also be thought of as perfect, intraspecific Batesian mimics (Brower et al., 1970). In contrast to Broom et al. (2005), Svennungsen and Holen (2007) found that populations with automimics can stabilize, leading to populations consisting of both defended and undefended individuals.

While these studies form a solid theoretical foundation for understanding the early evolution of defenses, a major group of organisms have been largely overlooked. These models assume defense operates at the individual level, where individuals with greater levels of defense are more likely to survive attack. This assumption does not apply to the common case of soft-bodied insects, such as aphids and caterpillars, for whom non-consumptive interactions with predators are still lethal. For these taxa, the effect of defense is frequency-dependent, where the effectiveness of the defense is affected by the proportion of defended individuals.

To address this gap in the current literature, we present two sets of game theoretical models. The first set uses classic, deterministic stable-strategy evolutionary game theory as described by Smith (1982). Here, instead of considering prey along a continuum, prey are treated as categorically defended or undefended. We used these models to ask how conditions such as predation pressure and the aversive effects of defense on predators affect the stability and equilibria of defended and undefended factions of populations when signals such as aposematic coloration are present vs. absent.

Like all models of this type, this first set assumes no variation in predator behavior, thus, parameters such as predator attack likelihood are constant. However, predators are known to alter their decision making based on internal physiological states, such as hunger (Barnett et al, 2007). Presumably, this would create a significant amount of stochasticity within an evolutionary system, which can shape selective pressures (Lenormand et al., 2009). As such, we create a second set of recursive, stochastic Wright-Fisher models (Tran et al., 2013), which use state-dependent, dynamic optimization (Clark and Mangel, 2008) of predator behavior as a means of selection. With these models we treat prey as either highly defended, moderately defended, or undefended, and investigate the role of prey population size, prey quality, and attack rates.

### Model Set 1: Deterministic Theory of Defense Games

#### Global Assumptions

These models assume population fitness, *W*, is fundamentally determined by two parameters, reproductive fitness, *V*, which represents the average reproductive rate of the population; and survivorship, *S*, which is the average proportion of individuals that survive to reproductive age, such that,

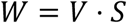

These models also assume prey use one of two mutually exclusive strategies, undefended or defended, where defended prey synthesize or sequester an invisible defense (e.g., chemical toxins) which negatively affects its predator. When a predator attacks, it kills prey, regardless of defense level. Thus, the advantage of defense is not to the individual, but comes from a predator’s learned experience and subsequent aversion to defended prey, which decreases a predator’s willingness to attack. These models also assume defense, whether synthesized or sequestered, has negative fitness consequences to prey, such as reduced growth and reproduction (Agrawal et al., 2021). Lastly, for simplicity, we assume all prey reproduce asexually.

These assumptions were analyzed using a typical payoff matrix (Tables 1 & 2), as described by Smith (1982), where a population consisting solely of defended prey has the payoff *E*(*D, D*) to each defended individual, while the payoff to defended prey in a mixed population of defended and undefended prey is *E*(*D, U*). Likewise, payoff to a pure undefended population is *E*(*U, U*) and for undefended prey in a mixed population the payoff is *E*(*U, D*). Payoffs for both prey types are mathematically similar to those of Leimar et al. (1986) and Broom et al. (2005), integrating intrinsic fitness, cost of defense, attack likelihood, and defense levels. Here, defended and undefended prey have the same intrinsic fitness, *v*, and defended prey suffer a fitness cost, *c*, for being defended. Intrinsic fitness, *v*, is always positive. The average cost of defense among defended individuals, *c*, reduces the intrinsic fitness proportional to the average level of defense *θ*, such that as *cθ* increases, fitness decreases. While Leimar et al. (1986) and Broom et al. (2005) set fitness to decrease exponentially with increasing levels of defense, we chose to do so linearly, such that

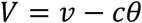

for reasons that will be apparent later.

**Table 1.** Payoff matrix, signals absent.

|  | <u>Defended</u> | <u>Undefended</u> |
| --- | --- | --- |
| <u>Defended</u> | $E(D, D) = (v - c\theta)(1 - \alpha e^{-\theta})$ | $E(D, U) = (v - c\theta)(1 - \alpha e^{-\theta P})$ |
| <u>Undefended</u> | $E(U, D) = v(1 - \alpha e^{-\theta P})$ | $E(U, U) = v(1 - \alpha)$ |

**Table 2.** Payoff matrix, signals present.

|  | <u>Defended</u> | <u>Undefended</u> |
| --- | --- | --- |
| <u>Defended</u> | $E(D, D) = (v - c\theta)(1 - \alpha e^{-\theta})$ | $E(D, U) = (v - c\theta)(1 - \alpha e^{-\theta P})$ |
| <u>Undefended</u> | $E(U, D) = v(1 - \alpha)$ | $E(U, U) = v(1 - \alpha)$ |

We assume the proportional cost of defense does not exceed intrinsic fitness such that fitness adjusted for the cost of defense is always greater than 0, meaning *cθ* < *v*. We also assume that in the absence of defense, attacks occur at an average constant per capita rate *α* such that survivorship,

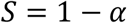

therefore, 0 < *α* < 1. We then multiply the fitness, adjusted for the cost of defense, by the likelihood of survival, to determine the realized fitness for defended prey:

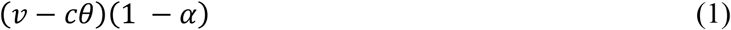

We, further, assume predator experience with defended prey decreases the prey’s likelihood of being attacked, *α*; and this influence of defense on *α* is greater than defense’s influence on intrinsic fitness. Therefore, like the models of Leimar et al. (1986) and Broom et al. (2005), decrease in the likelihood of being attacked declines exponentially with respect to defense, *θ*, such that,

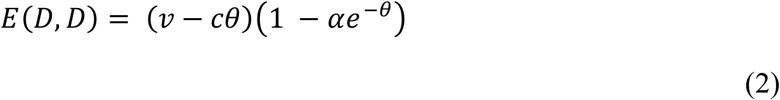

In contrast, undefended prey experience no cost in defense and in pure populations predators experience no aversion, ergo:

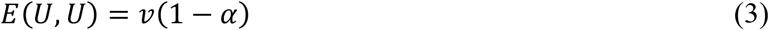

See tables 1 and 2.

In theory, both *c* and *θ* could take on negative values, and there may be empirical cases in which *c* does so. For example, *Uthesia* spp. use pyrrolizidine alkaloids (PA’s) as a means of defense; but males also convert PA’s into dihydropyrrolizine hydroxydanaidal which is a necessary courtship pheromone (Conner et al., 1981; Conner and Weller, 2004). Arguably, in this scenario *c* would be negative, since increases in *θ* not only contribute to likelihood of survival but also increases fitness—at least for males. However, for our purposes, *θ* must necessarily be positive for two reasons. First, if *θ* were negative and *c* was positive, defense would increase the adjusted intrinsic fitness despite there being a cost to defense. Second, values of *θ* less than 0 would mean the likelihood of attack increases with increasing levels of defense, meaning “defended” individuals are not truly defended. Thus, we assume the condition that *θ* ≥ 0.

#### Signals Absent

When predators are unable to distinguish defended prey from undefended prey (i.e., signals are absent; Table 1), in mixed populations the likelihood of defended prey being attacked is greater than that of a purely defended population. This is because the effect of *θ* decreases as the proportion of defended prey, *P*, decreases due to the positive reinforcement predators receive from attacking undefended prey. Thus, the payoff for defended prey in a mixed population is,

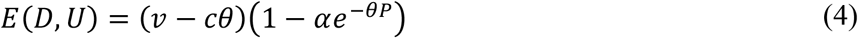

In contrast, for undefended prey in mixed populations, the likelihood of being attacked is less than that of a purely undefended population, as predators associate undefended prey with defenses from defended members of the population. Thus, the effect *θ* has on the likelihood of being attack, *α*, is included in the payoff for undefended prey, with its effects increasing as the proportion of defended prey, *P*, increases, but not with respect to intrinsic fitness, such that,

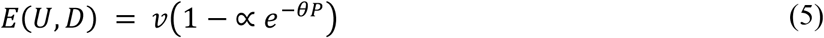

#### Signals Present

When signals are present (Table 2), we assume predators can perfectly identify defended prey. As such, intuition might suggest when signals are present the payoff for defended prey in an admixed population (*E*_*S*_(*D, U*)) is the same as an exclusively defended population (equation 2). However, because the effect of defense (*θ*) is dependent on predator learning, it is still frequency-dependent, such that the effect of *θ* decreases as the proportion of prey that are defended (*P*) approaches 0. Therefore, when signals are present, realized fitness for defended prey in mixed populations, *E*_*S*_(*D, U*), is approximately equal to defended prey in an admixed population where signals are absent (equation 4). In contrast, for undefended prey in mixed populations, realized fitness is dependent on whether signals are present vs. absent. As mentioned above, when signals are absent (Table 1), undefended individuals in mixed populations experience fewer attacks as the number of defended individuals increases and experience no loss to intrinsic fitness. However, when signals are present we assume predators can distinguish defended prey from undefended prey with 100% accuracy, thus the effect of *θ* on undefended prey is removed. As such, when signals are present,

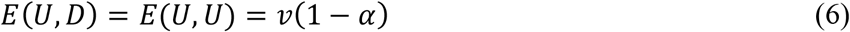

Given these assumptions, when signals are absent the corresponding payoff matrix is:

And when signals are present:

Using these models, we can now analyze the evolutionary stability of either strategy, as well as mixed strategies, when signals are present vs. absent.

#### Analytical Analysis

##### Undefended Populations

The undefended strategy is evolutionarily stable (i.e., the defended strategy cannot invade) when the payoff to being an undefended individual in a purely undefended population is greater than the payoff of being a defended individual in a mixed population of defended and undefended individuals. Therefore, when *E*(*U, U*) > *E*(*D, U*), the undefended strategy is evolutionarily stable. This means when signals are absent the undefended strategy is evolutionarily stable when,

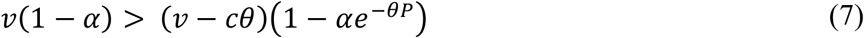

This can be arranged such that,

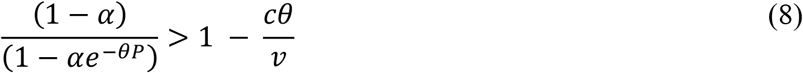

This itself is not very insightful, however if we assume certain boundary conditions such as defense has no cost to defended prey (i.e., *c* = 0) the inequality simplifies to,

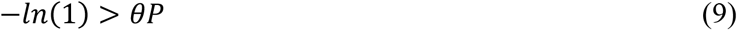

which, given conditions such as *P* is a proportion and must be between 0 and 1, and *θ* is positive, the inequality is false, as *θP* is necessarily positive. Since this inequality is always false, we can conclude that the undefended strategy is evolutionarily unstable when there is no cost to prey adopting a defended strategy. Likewise, if there are no predators attacking this population of prey, such that *∝* = 0, undefended is always evolutionarily stable given defense has a non-zero cost, as the inequality would simplify to 0 < *c*, which is always true given the previously stated conditions. The same trivial scenarios are true when signals are present as well.

##### Defended Populations

The defended strategy is evolutionarily stable (i.e., the undefended strategy cannot invade) when the payoff of being a defended individual in a purely defended population is greater than the payoff of being an undefended individual in a mixed population. *E*(*D, D*) > *E*(*U, D*). Therefore, when signals are absent the defended strategy is stable when,

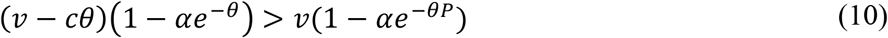

This simplifies to,

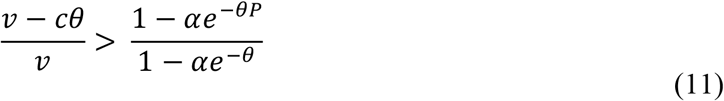

When the cost of defense is 0, *c* = 0, 1 − *αe*^−*θ*^ > 1 − *αe*^−*θP*^, which is true for all values of *P* < 1, the defended strategy is evolutionarily stable. However, when predators are absent, *∝* = 0, equation 10 simplifies to,

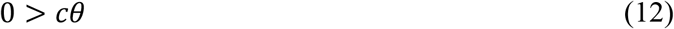

which is always false given the assumption *c* and *θ* are always positive. Thus, the defended strategy can be invaded by the undefended strategy. Both of these boundary condition, both *c* = 0 and *α* = 0, outcomes make intuitive sense and are also same when signals are present.

#### Numerical Analysis

##### Mixed Populations

The most interesting solutions, of course, are the non-trivial cases. However, because of the relative complexity of these models, general analytical solutions are largely intractable. Therefore, it is useful to impose biologically plausible constraints on the models and solve the equations numerically.

We focused our analyses to address whether defended prey would increase in a population, thus we computed the change in relative fitness for the defended strategy, given a set of plausible assumptions. We calculated the mean change in relative fitness for defended prey according to Smith (1982) where fitness, *W*, of the defended phenotype is,

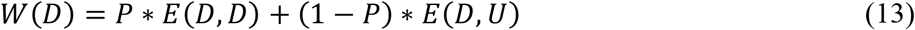

and fitness of the undefended phenotype is,

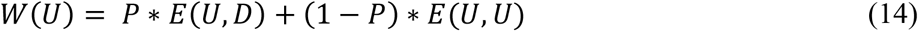

Since we assumed prey reproduce asexually, prey reproduce in proportion to their fitness. Thus, we can find the proportion of defended prey in the next generation, *P*_*t*+1_ as:

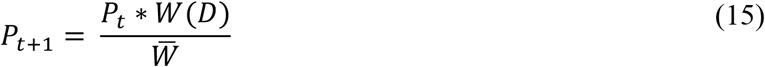

where,

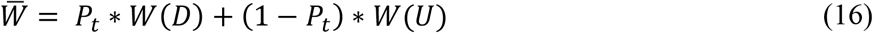

Thus, the change in proportion of defended prey, 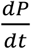, is:

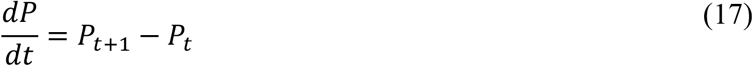

To simplify the numerical solutions, we assumed the proportional cost of defense to intrinsic fitness is exactly equal to the level of defense, such that *c* = 1. Therefore, *E*(*D, D*) and *E*(*D, U*) are equal to,

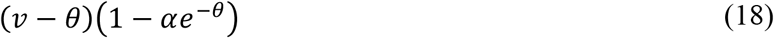

and

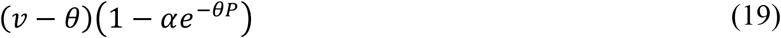

respectively.

We established some parameter bounds and values based on basic biological principles and for heuristic comparison. We chose to limit defense such that 0 < *θ* < 3. We chose 3 as the upper limit because when the likelihood of being attacked is 100% in the absence of defense (*α* = 1), at *θ* = 3, approximately 5% of the population would still be attacked. This seemed a reasonable assumption as naïve predators would still need to attack some portion of a purely defended prey population to learn to avoid said prey. We assumed *v* = 8 so that a population would experience no more than a 40% loss in fitness when maximally defended. This may be a conservative estimate, as monarch caterpillars can experience greater than 80% reductions in growth over an 8-day period when maximumly defended (Agrawal et al., 2021). However, we are less concerned with exact values and are more interested in the qualitative predictions of these models. While investigating the effect of *∝*, we chose to set *θ* = 1.5, as it is the midpoint for the range of *θ* values. While investigating the effects of *θ*, we set *∝* = 0.3. Just as with *v*, these values are arbitrary, as we are interested in the qualitative features of these models.

##### Evaluation

For single parameters we evaluated change in defended-prey frequency across a grid of our parameter of interest against initial defended-prey frequencies (*P*). We visualized the direction and magnitude of this change according to *dP*/*dt*, such that values > 0 represent increases in defended-prey frequencies and are represented in red, while values < 0 represent decreases in defended-prey frequencies, represented in blue. Magnitude was calculated as the absolute value of *dP*/*dt*. As magnitude decreased, grid points became lighter, with values at or near 0 being white, representing the null-clines, where population composition does not change between generations.

### Model Set 2: Stochastic Models

The first model set focused almost entirely on prey, treating predator-dependent parameters (i.e., attack rates and effect of defense) as average, unvarying constants. There, predators are passive participants. In contrast, our second model set explicitly accounts for predator choice, fitness, and includes stochastic effects, which here arise due to variation in predator initial states and predators behaving in ways that maximize their own future fitness. This top-down paradigm results in predicted attack rates and illustrates the effect of defense being more dynamic and varying between replicates and iterations. To construct these stochastic games, we identified the optimal predator behaviors (those that maximized predator fitness) using dynamic optimization modeling (Clark and Mangel, 2008). Like the previous model set, one model represents the case where signals are present, while the other represents when signals are absent. We then simulated predators foraging in an environment with a fixed number of prey that varied in levels of defense. Predators that are encountering prey in the environment can attack and consume them, skip attacking and continue to forage within the current patch, or leave the foraging patch in search of other prey types. Of course, in the signals absent model they do not know what type of prey they have encountered, however their behavior is based on their expectation and the consequences for their fitness (Clark and Mangel, 2008). In the case where signals are present, of course that knowledge influences decisions directly. Prey that do not experience predation, reproduce at the end of each iteration, and thus the distribution of prey types in the next generation reflects changes in the proportion of defended prey as a result of survivorship as well as variation in fecundity, which decreasing as defense increases (as in our previous models).

#### Assumptions

We chose a spatially implicit, discrete, multi-generational model, where predators operate across a range of energy states, *x*, from *x*_*crit*_ to *x*_*max*_. Here, predators forage for a fixed amount of time, known as the time horizon, *T*_*max*_, which is determined by the number of available prey in the environment, *K*, and the maximum proportion of those prey that can be found, *A*, so that,

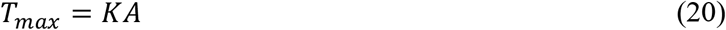

In effect, *K* represents the prey population size, and *A* represents the maximum attack rate, which reflects a multitude of different factors including predator mobility and prey detectability. At each time step, *t*, predators are exposed to one of three prey types *Y*_*j*_, where *j* belongs to a set of three variables, *j* ∈ {1,2,3}, and *Y*_1_ = *Undefended, Y*_2_ = *Moderately Defended*, and *Y*_3_ = *Highly Defended*. We chose three different prey types as a matter of convenience, but the types could be further subdivided an arbitrary number of times (e.g., *Highly Palatable, Undefended, Neutrally Defended, Highly Defended, Extremely Defended*). Regardless, 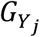 represents the amount of energy gained or lost by a predator when it consumes prey of type *Y*_*j*_, which in this model is defined by the maximum prey quality, *Q*. For this model we assume undefended prey represent the highest quality prey, where the predator energy state increases by

*Q*, where *Q* can be any positive, even integer greater than 2 such that,

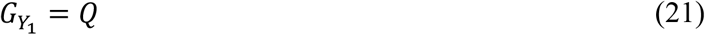

Moderately defended prey are of neutral value, meaning prey are defended enough that predators do not gain energy but see no loss in energy due to foraging and metabolic costs. As such, predators feeding on moderately defended prey experience a net 0 change in energy. Thus,

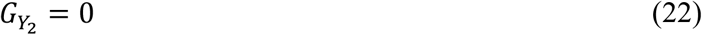

Finally, highly defended prey are of such poor quality they produce a net loss of energy, such that,

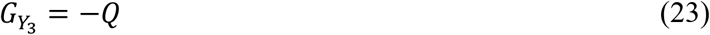

We assume this loss of energy is due to emetic effects. We chose to do so as vomiting is a common predator reaction to some chemically defended prey (Brower, 1969). Likewise, vomiting would lead to an immediate loss of stored energy, simplifying energetic losses in the model to a single time step.

In each time step, predators encounter a prey item and can choose one of three strategies: consume the prey, avoid the prey and continue to forage within the focal environment, or leave the environment for a new one with other prey. If predators choose to continue to forage inside the focal environment, they necessarily lose one unit of energy due to the metabolic costs of maintenance and movement.

For simplicity, we assume predators do not use Bayesian updating within a generation, meaning optimal strategies are determined by the probability of encountering each prey type at the start of each generation, and do not change dynamically until the prey population reproduces. This assumption allows us to isolate the effects of state-dependent predator decision-making.

Because of the ability to optimize their behavior, predators act within the model with a certain degree of agency. Prey, however, do not.

Like the previous model set, we assume prey that have been attacked by predators always die and defenses are invisible, meaning that unless signals are present, predators cannot discern between defended and undefended individuals within a population. Similarly, we assume defense carries a cost to intrinsic fitness such that each surviving individual of prey type *Y*_*j*_ reproduces *N*_*j*_ individuals. To simplify the model and because both variables are a function of energy, we assume that the benefit to the predator of the prey, *Q*, is likewise a component of prey fitness, such that each undefended individual produces 200 + *Q* offspring per generation, moderately defended individuals produce 200 offspring, and highly defended individuals produce 200 − *Q* offspring. Depending on the value of *Q*, this provides undefended individuals a minimum 1.9% and a maximum 5.8% greater fecundity than any defended prey. Because this is a discrete model, as with *K, Q* is always an integer.

#### Dynamic optimization model

To determine the time and state dependent optimal behaviors of predators, we used a dynamic programming algorithm as described by Clark and Mangel (2000). Here future predator fitness is dependent on their energy state at the end of the model, determined by a general terminal fitness function. In defining our terminal fitness function, we assumed predators were small insectivorous, passerine birds (a common predator of soft-bodied insects). Passerines often have relatively high metabolic rates (McNab, 2016), suggesting high prioritization of immediate tradeoffs. As such we chose a fitness coefficient with respect to energy of 0.5. Likewise, stored energy would have few, if any diminishing returns. Therefore, we chose to make our terminal fitness equation linear, such that,

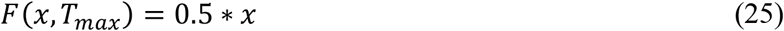

where *x* is the energy state of the predator.

#### Signals absent

When signals are absent, energy is the only state variable that influences predator behavior; thus,

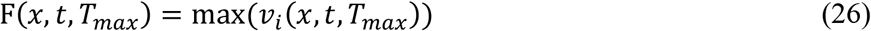

where *v*_*i*_ is the expected future fitness of strategy *i*, where *i* belongs to the set of behaviors *I* and *I* = {*Consume Prey, Forage Within System, Forage Outside System*}.

Here, *v*_1_ (consume prey) is influenced by the proportion of each prey type, *p*(*Y*_*j*_). Because *p*(*Y*_*j*_) is a proportion, relative to the total number of prey available it is necessarily the case that the proportions sum to one. That is,

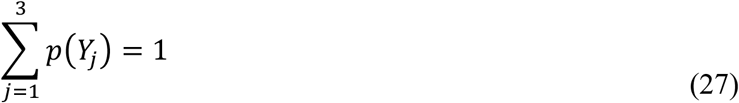

This means the expected future fitness at time *t* is simply the sum of each proportion of prey type multiplied by the expected future fitness moving one time-unit forward, given the new energy state after consuming that prey type, *x*′,. Ergo,

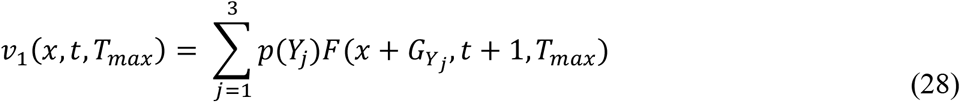

For *v*_2_ future energy, *x*′, is equal to *x* minus metabolic and movement costs, which in our model is one unit of energy. Therefore,

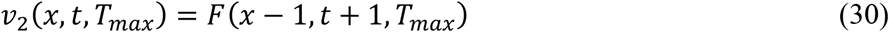

For predators foraging outside the local system, we used a time-dependent but state-independent function to represent future fitness such that,

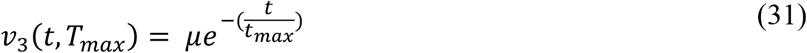

where *µ* is the average probability of encountering prey in a neighboring environment, which decays exponentially with time.

##### Signals present

In the case where signals are present the terminal fitness function is the same as equation 25. However, the expected future fitness of strategy *i* includes the prey type *Y* as a state variable, such that,

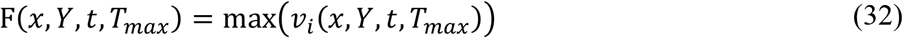

While *Y* is a known state at time *t*, that state changes through time as they encounter prey of different types, thus our fitness equations for eating and foraging must account for the future probabilities of encountering each prey type. Thus,

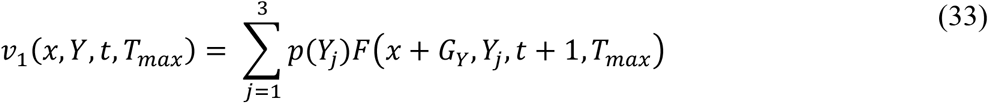

and,

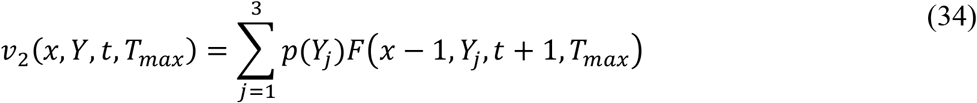

while the equation for foraging in a neighboring patch remains the same as the signals absent model.

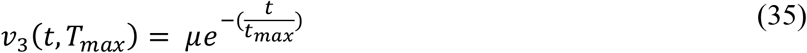

#### Forward simulation and selection

We began each replicated simulation of prey being attacked by predators using the optimal foraging strategy at generation 1, with *K* prey individuals. Since we were largely interested in how an undefended population shifts to a defended population, at generation 1 the initial population of *K* individuals was randomly sampled using a weighted probability, with a 75% chance any one individual was undefended, 15% chance they were moderately defended, and a 10% chance they were highly defended. Our models did not include mutation. From that population we calculated the proportion of each prey type, to determine the probability of a predator randomly encountering each prey type, *p*(*Y*_*j*_), which was used to generate the dynamic optimization model. We then randomly assigned a single predator an energy state, *x*, between *x*_*crit*_ and *x*_*max*_, discrete and uniformly distributed. Moving forward through time within that first generation, we sampled which prey type the predator encountered, with the probability of encountering each prey type weighted according to *p*(*Y*_*j*_). The predator then utilized the optimal strategy according to its state(s).

If the predator consumed the prey (*i* = 1), predator energy, *x*, changed for the next time step to *x*′, bounded by *x*_*max*_ and *x*_*crit*_ where,

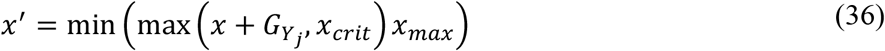

and an individual of prey type *Y*_*j*_ was removed from the population.

If the predator continued foraging within the system, energy *x* changed to *x*′, where,

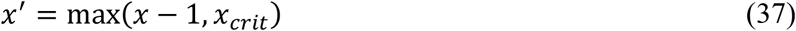

If the initial predator foraged outside of the system, or if *x*^′^ = *x*_*crit*_ suggesting the predator died, this created a space for a new predator with a new random initial energy, as above, to forage starting at the next within-generation time step. This process continued until *t* = *T*_*max*_, at which point we implemented the fecundity equation on all surviving prey individuals. Once all survivors reproduced, we randomly selected *K* individuals from that cohort to represent the new population of prey for the next generation. Again, we determined *p*(*Y*_*j*_), this time based on this new cohort, and implemented the dynamic optimization model, repeating simulations until they reached 50 generations. We replicated this procedure 1000 times for each combination of parameters (*A, K*, and *Q*), which consisted of one low value and one high value for each parameter (Table 3), and each model type. We evaluated our simulations according to prey composition at the 50^th^ generation, observing the proportion of replicates in which all prey were of one type (Highly Defended, Moderately Defended, Undefended) or some admixture of prey types. We then conducted sensitivity analysis on the parameters *A, K*, and *Q*, to see the general effect each parameter had on the composition of admixture populations using LOESS smoothing of our simulations. We fixed *A* = 0.4 for evaluations of *K*(25: 200) and *Q* (2: 8), *K* = 50 for *A* (0.2: 0.8) and *Q*, and *Q* = 0.6 for *A* and *K*.

**Table 3.**
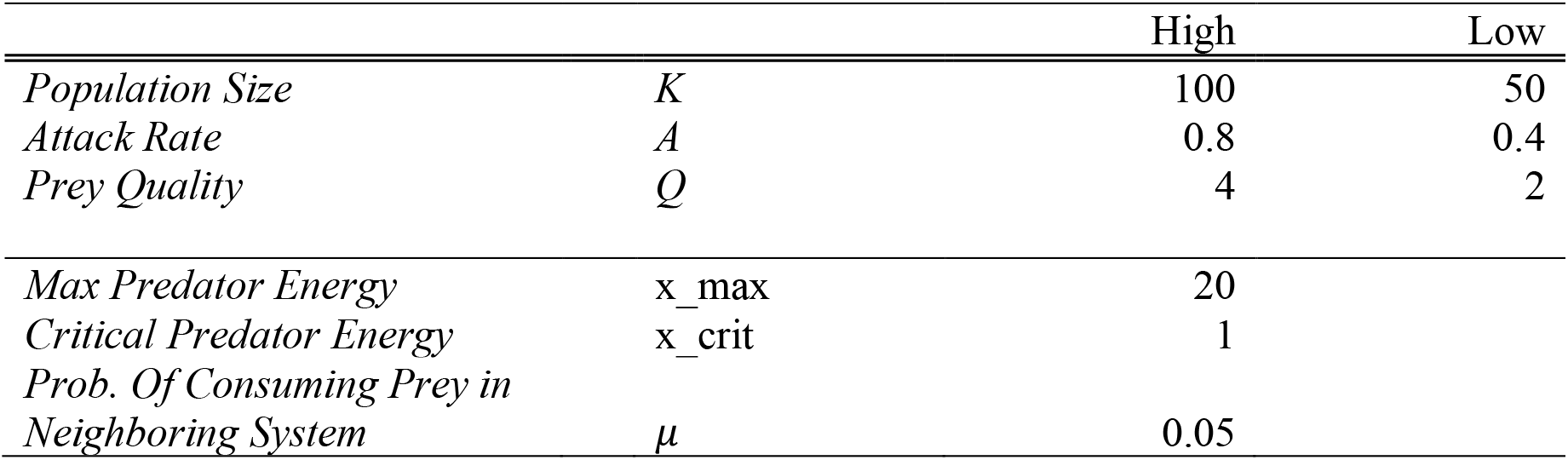
Parameters for the second model set.

All models and simulations were written in C++ then implemented in R (4.5) using the Rcpp package (1.1). Figures for model set 1 were generated using base R graphics, while figures for model set 2 were generated via the ggplot2 package (3.5.2) using an R studio (2025.5.1.513) interface.

## Results

### Model Set 1. Deterministic Theory of Defense Games

We evaluated whether defended and undefended phenotypes were evolutionarily stable, whether mixed-population equilibria were stable, and how the composition of those equilibria varied with attack probability and defense level, given the presence or absence of signals. This first model set revealed several qualitative features. First, for both models, with signals present and absent, the undefended phenotype is stable below some threshold likelihood of being attacked, *α* (Figures 1A and 1B), meaning the defended phenotype cannot invade a population if the likelihood of being attacked is too low. Unsurprisingly, the actual threshold is dependent upon other parameters. For example, increasing intrinsic fitness, *v*, decreases this attack threshold, while increasing defense and its effects on predator behavior, *θ*, increases the attack threshold (Figure 2A). In contrast, the undefended phenotype is stable above some defense threshold, *θ*. That threshold also increases with increasing intrinsic fitness, *v*, and the likelihood of being attacked, *α*, (Figure 2B). Notably, the initial downward curvature of the zero-change lines in Figure 1A and 1B and the lower parameter space dominated by decreasing values of *dP*/*dt* show that as attack probabilities, *α*, decrease, the initial proportion of undefended prey necessary to avoid invasion by defended prey decreases as well. In contrast, defense, *θ*, appears to have the opposite effect, with increasing levels of defense requiring lower initial undefended prey frequencies to avoid invasion by defended prey.

**Figure 1.**
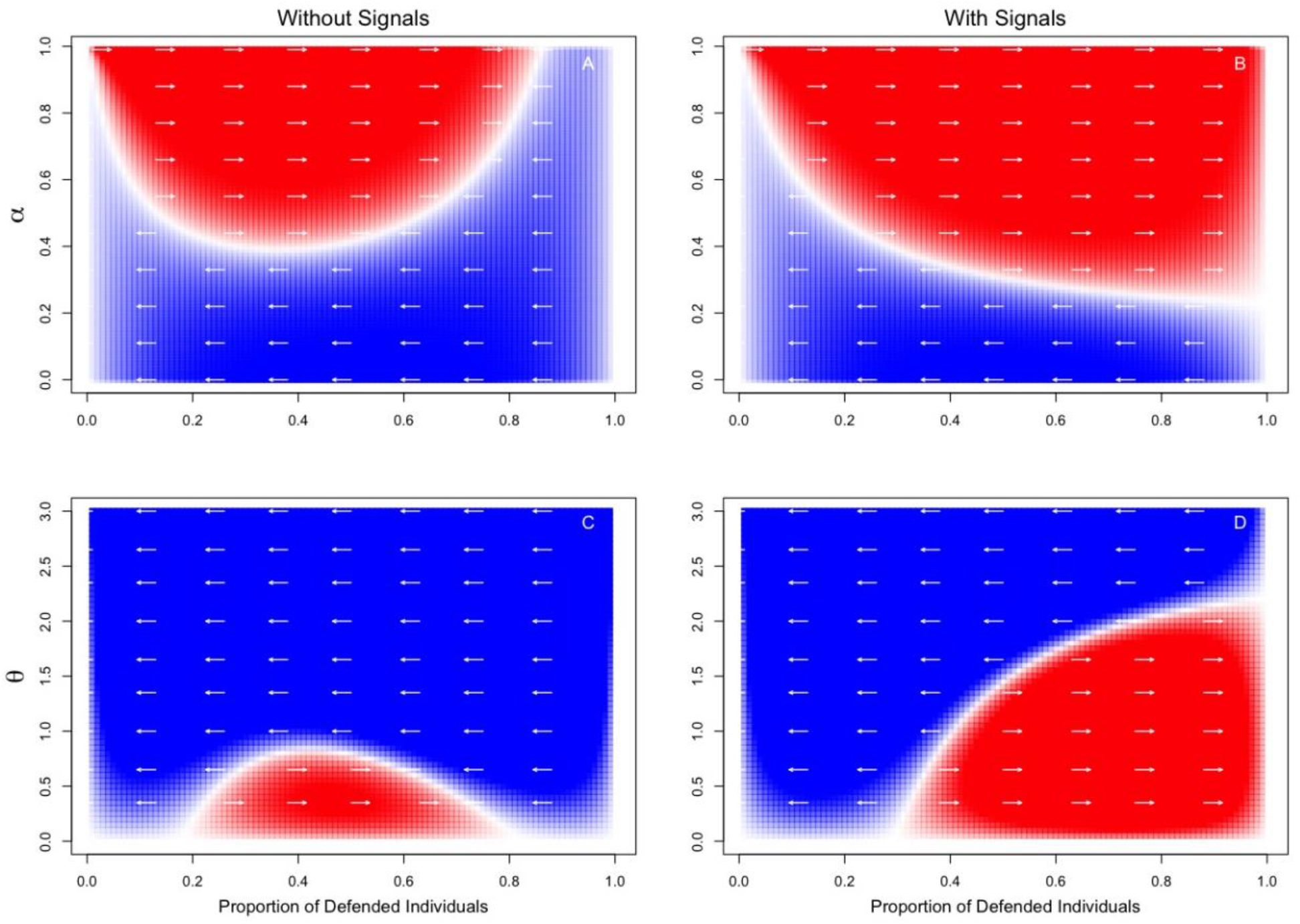
Stability dynamics of model set 1. for different levels of *α* (likelihood of being attacked) for the model without signals A) and the model with signals B); and different levels of *θ* (level of defense) for the model without signals C) and the model with signals D). Red indicates the proportion of defended individuals are increasing, blue indicates proportion of defended individuals are decreasing, white represents the null-clines.

**Figure 2.**
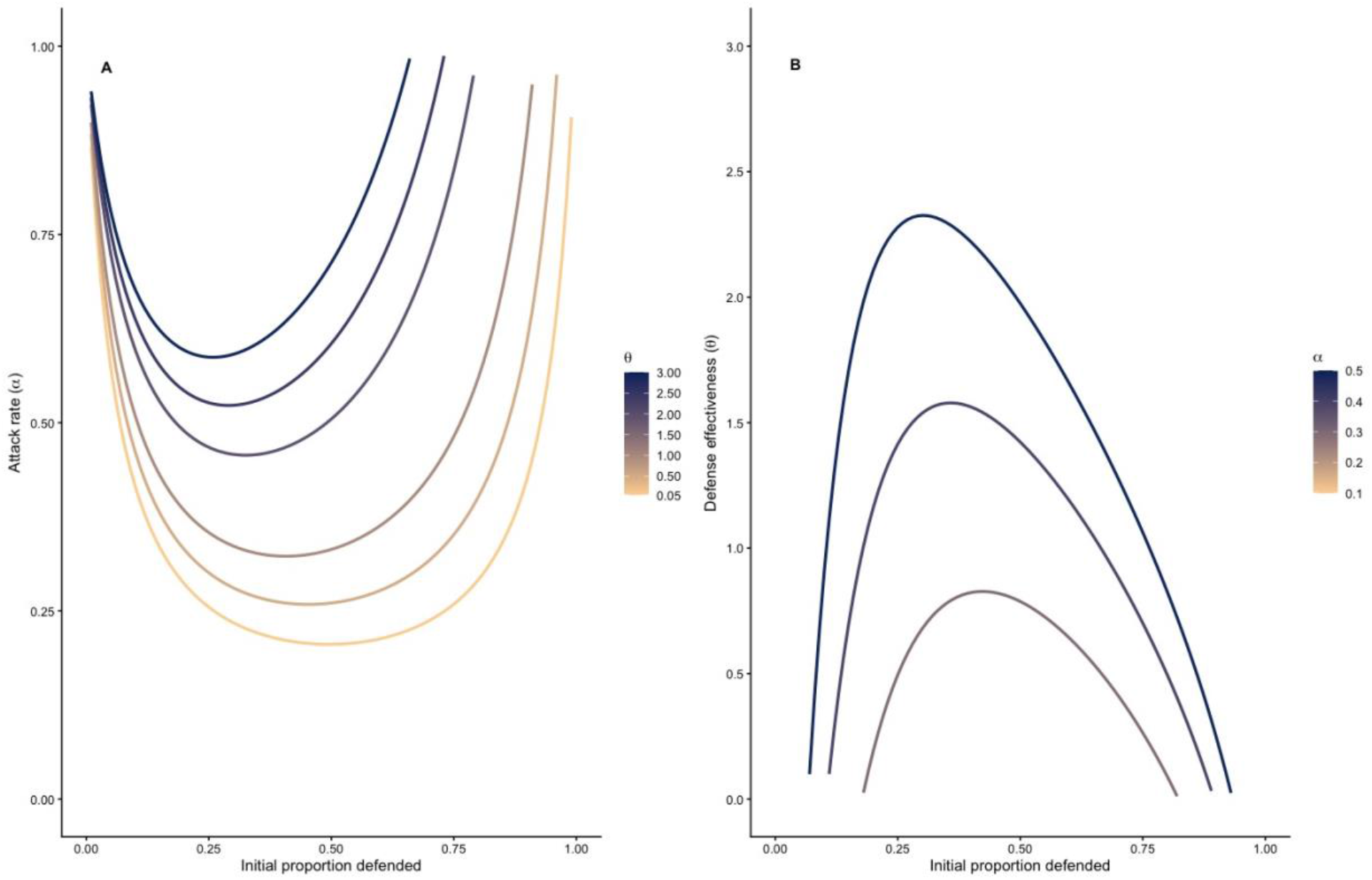
Null-clines for attack likelihood (*α*) against initial proportion of defended prey with increasing values of defense effectiveness (*θ*) A., and *θ* against proportion of defended prey for increasing values of *α* B.

**The signals present model** showed another notable feature. Specifically, equilibria across all mixed populations are unstable, meaning the population will eventually deviate from a purely defended or undefended population, depending on the parameters and the initial proportions of each phenotype. For example, the defended phenotype was able to invade and remain stable when the likelihood of being attacked, *α*, was high. The threshold attack probability required for defended phenotype to invade decreased as the initial proportion of defended prey, *P*, increased, indicating the initial frequency of defended prey affected the conditions under which defense could become established. However, the defense threshold *θ* leading to stability of the defended phenotype increases as the initial proportion of defended prey, *P*, increases. In contrast, when **signals are absent**, most equilibria are stable and the defended phenotype appears incapable of complete invasion, meaning there are always at least some undefended individuals mixed within the population. Moreover, there appear to be bifurcation-like points, thresholds where the mixed population equilibria are unstable at lower initial defended prey frequencies, stabilizing at higher initial frequencies. To understand the effects of other parameters on these points, we conducted a complimentary analysis of the signals-absent model to identify the effects of one parameter on the zero-change conditions of another. For each parameter combination we evaluated the model over 500 evenly spaced values of the causal parameter and identified the intervals in which consecutive evaluations of *dP*/*dt* had opposite signs (positive vs. negative), estimating the first zero crossing using the uniroot function in R, with local minima and maxima representing the bifurcation-like threshold.

Subsequently, we found that bifurcation-like thresholds appear to increase with respect to attack probabilities (*α*) as defense (*θ*) increases (Figure 2A). Likewise, thresholds with respect to defense (*θ*) increase as attack probabilities (*α*) increase (Figure 2B).

### Model Set 2. Stochastic Models

Like Model Set 1, Model Set 2 was designed to examine how prey quality and predation pressure affect the stability and composition of prey populations. However, in Model Set 2, we explicitly accounted for prey population size and selection imposed due to predator state-dependent behavior. We assumed a particular defense type was fixed when 100% of a population consisted of that phenotype after 50 generations. Similar to Model Set 1, admixtures were almost entirely absent from the **signals present model**, the highest proportion being 12 out of 1,000 independent simulations (*K* = 25, *A* = 0.4, and *Q* = 6; Table 5). Likewise, across all parameter combinations the majority of simulated populations that did end with a fixed phenotype after 50 generations (i.e., were stable) were overwhelming in favor of the highly defended phenotype, ranging from 76.5% to 99.3% (Table 5). Small populations (*K* = 25) with low attack rates (*A* = 0.4), had the lowest proportion of independent simulations in favor of the highly defended phenotype, with 76.5% of the simulations being completely invaded by the highly defended phenotype after 50 generations. In contrast, in the **signals absent model**, for all parameter combinations the undefended prey type was fixed in the vast majority of cases. However, when population size was small (K=25) fixed populations for the highly defended and moderately defended prey types did occur, although this was infrequent (2.6-6.2% and 9.8-15.1%, respectively). At higher population size (K=100) fixed populations for moderately defended prey also occurred. Admixed populations did occur for some combinations (e.g., large populations, and lower attack rates), occurring in relatively high frequency (37% and 25%; Table 4).

**Table 4.** Proportion of final populations Signals Absent model.

| **K** | **A** | **Q** | **Highly**<br>**Defended** | **Moderately**<br>**Defended** | **Undefended** | **Mixed** |
| --- | --- | --- | --- | --- | --- | --- |
| 25 | 0.4 | 4 | 0.042 | 0.098 | **0.813** | 0.047 |
| 25 | 0.4 | 6 | 0.026 | 0.106 | **0.823** | 0.045 |
| 25 | 0.8 | 4 | 0.062 | 0.151 | **0.778** | 0.009 |
| 25 | 0.8 | 6 | 0.040 | 0.149 | **0.802** | 0.009 |
| 100 | 0.4 | 4 | 0.000 | 0.003 | **0.626** | **0.371** |
| 100 | 0.4 | 6 | 0.000 | 0.003 | **0.747** | **0.250** |
| 100 | 0.8 | 4 | 0.000 | 0.065 | **0.742** | 0.193 |
| 100 | 0.8 | 6 | 0.000 | 0.069 | **0.800** | 0.131 |

**Table 5.** Proportion of final populations Signals Present model.

| K | A | Q | Highly<br>Defended | Moderately<br>Defended | Undefended | Mixed |
| --- | --- | --- | --- | --- | --- | --- |
| 25 | 0.4 | 4 | **0.765** | 0.028 | 0.199 | 0.008 |
| 25 | 0.4 | 6 | **0.768** | 0.018 | 0.202 | 0.012 |
| 25 | 0.8 | 4 | **0.912** | 0.011 | 0.077 | 0.000 |
| 25 | 0.8 | 6 | **0.932** | 0.012 | 0.056 | 0.000 |
| 100 | 0.4 | 4 | **0.988** | 0.000 | 0.003 | 0.009 |
| 100 | 0.4 | 6 | **0.980** | 0.000 | 0.011 | 0.009 |
| 100 | 0.8 | 4 | **0.993** | 0.001 | 0.005 | 0.001 |
| 100 | 0.8 | 6 | **0.987** | 0.001 | 0.007 | 0.005 |

Because there were so few admixture populations in the signals present model, we limited our sensitivity analyses to the **signals absent model**. Average final prey type proportions showed undefended prey were the dominant phenotype at the lowest attack rates, with moderately defended prey dominating around *A* = 0.4 then quickly declining as attack rates increased. The highly defended phenotype increased as the dominant phenotype exponentially with increasing attack rates (*A*, Figure 3A). In contrast, the undefended prey type remained the dominant phenotype as population size (*K*) increased (Figure 3B), increasing in mean proportion, with similar patterns for increasing levels of *Q* (Figure 3C).

**Figure 3.**
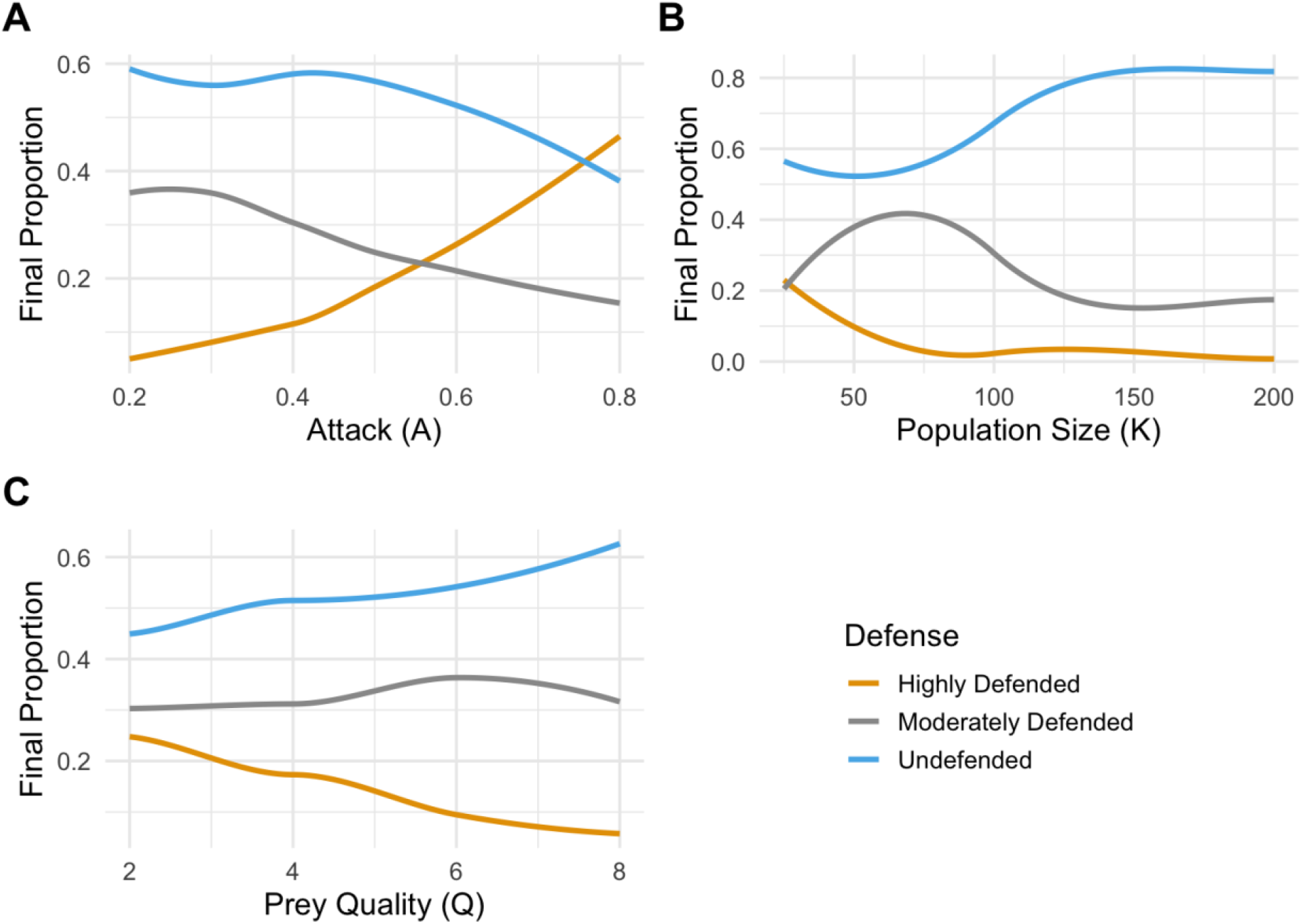
Sensitivity analysis for final proportions of mixed populations from model set 2. for different levels of *A* panel A., *K* panel B., and *Q* panel C.

## Discussion

Despite the importance of defense to fitness and survival of prey, undefended phenotypes are surprisingly likely to persist according to both model sets. Broom et al. (2005) showed that low predation pressure can make investment in secondary defenses disadvantageous when defenses are invisible. According to our first model set, low predation pressure appears to diminish the selective advantage of defense regardless of whether signals are present or absent. However, as attack probabilities increase, defense becomes more advantageous, leading to complete invasion when signals are present and admixed populations when signals are absent.

The negative effect low attack probabilities have on invasion by the defended phenotype is likely attributable to the cost of defense, again consistent with Broom et al. (2005), as the initial proportion of undefended individuals necessary to maintain a purely undefended population decreases as the level of defense increases among the defended individuals of a population, which is exacerbated for systems in which signals are absent. This yields a counterintuitive prediction with respect to automimicry. Specifically, if we define a stable automimic system as one in which signals for determining which prey are defended vs. undefended are absent, and there exists stable equilibria of mixed populations, then the signals absent model appears to be a stable automimic system given three conditions. Specifically, predation pressure must be relatively high, defense levels among defended individuals must be relatively low, and there must be a minimum initial proportion of defended individuals greater than the bifurcation-like threshold. The last of these conditions itself makes intuitive sense, but also suggests implications for the inclusion of the defended phenotype into a population of undefended individuals.

Explicitly, the initial proportion of defended individuals must be relatively large. One way in which this may manifest is through smaller prey populations, which *de facto* would increase the initial proportion of defended individuals. These findings are consistent with other theoretical predictions that prey density can influence the evolutionary benefits of defense, although the underlying mechanisms differ. Using a functional-response model, Jeschke (2006) demonstrated that the benefits of defense depend on both prey density and the stage of the predation sequence affected by the defense. Defenses that affect predator encounter, detection, attack, or consumption were predicted to be most beneficial at low prey densities, whereas defenses that affect searching were predicted to be beneficial at intermediate densities. Our results expand upon this density-dependent perspective, demonstrating prey population size can also influence invasion by defended phenotypes through its effects on their initial frequency. In particular, when defended and undefended prey are indistinguishable, smaller populations were more conducive to invasion by the defended phenotypes.

Other ways in which the initial proportion of defended prey could be relatively large could be due to clutch site effects, predator mobility, and maternal behavior. As an example, female butterflies that sequester defenses from host plants may choose to oviposit a disproportionate amount of eggs on host plants with higher concentrations of those sequesterable defenses (Lefevre et al., 2012). Alternatively, females may lay a disproportionate amount of eggs on low toxicity host plants, leading to clusters of defended and undefended individuals. For an avian predator this heterogeneity may be inconsequential, but for smaller, invertebrate predators with restricted ranges, a clutch may constitute a substantial fraction of the prey type experienced by the predator.

Conditions leading to stability of defended and undefended prey in the second model set follow similar patterns to the first. Namely, greater predation pressure significantly affects fixation by highly defended prey as well as dominance of highly defended prey among mixed populations. While prey population size is inferred to affect the invasion by defended individuals in the first model set, the second model set addresses this directly. For the signals present model, larger population sizes increase the proportion of outcomes in which highly defended individuals fully invaded the population. When signals are absent, however, larger populations had the opposite effect, congruent with the first model set. That is, when signals are absent and prey population sizes are small, defended individuals are more likely to invade undefended populations, whereas population size has little effect when signals are present and attack rates are high. This is not to say population size has no effect on the maintenance of defended phenotypes and only speaks to the ability of defended individuals to invade. In fact, defended prey that adopt conspicuous warning coloration are often found to be gregarious (Gamberale and Tullberg, 1998).

Apart from their qualitative predictions these models are also of practical value to empiricists, particularly model set 1. As mentioned above, the signals absent model from model set 1 provides a deterministic function for investigating populations that display automimicry. By simply collecting data on the proportion of automimics to models in a population over time, it would be possible to obtain estimates on a natural population’s intrinsic fitness, cost of defense, intrinsic predator attack probabilities, and the effects of defense on predator attacks. Moreover, both model sets provide robust, theoretically informed models that can be statistically evaluated to infer whether a particular prey feature, or feature of their environment, is being used as a signal for prey defense (Martin et al., *under review*).

## Acknowledgments and Contributions

We would like to thank Dr.’s Heather McAuslane, Jaret Daniels, and Phillip Hahn for their helpful feedback on an earlier version of this manuscript. This material is based upon work by CStM supported while serving at the National Science Foundation. The research questions, theoretical framework, and models presented were developed collaboratively by both authors.

